# Genome-wide DNA methylation profiles in smoking discordant and concordant monozygotic twin pairs

**DOI:** 10.1101/2022.08.17.504357

**Authors:** Jenny van Dongen, Gonneke Willemsen, BIOS Consortium, Eco J.C. de Geus, Dorret I. Boomsma, Michael C. Neale

**Affiliations:** Department of Biological Psychology, Vrije Universiteit Amsterdam, Amsterdam, The Netherlands; Amsterdam Public Health Research Institute, Amsterdam, The Netherlands; Amsterdam Reproduction and Development (AR&D) Research Institute, Amsterdam, The Netherlands; Biobank-based Integrative Omics Study Consortium; Virginia Institute for Psychiatric and Behavioral Genetics, Virginia Commonwealth University, Richmond, USA

**Keywords:** Cigarettes, EWAS, twin study, addiction, causality, epigenetic, smoke, Netherlands Twin Register

## Abstract

Smoking-associated DNA methylation levels identified through epigenome-wide association studies (EWAS) are generally ascribed to smoking-reactive mechanisms, but the contribution of a shared genetic predisposition to smoking and DNA methylation levels is typically not accounted for. We exploited a strong within-family design, i.e., the discordant monozygotic twin design, to study reactiveness of DNA methylation in blood cells to smoking and reversibility of methylation patterns upon quitting smoking. Illumina HumanMethylation450 BeadChip data were available for 769 monozygotic twin pairs (mean age=36 years,range=18-78, 70% female), including pairs discordant or concordant for current or former smoking. In pairs discordant for current smoking, 13 differentially methylated CpGs were found between current smoking twins and their genetically identical co-twin who never smoked. Top sites include multiple CpGs in *CACNA1D and GNG12*, which encode subunits of a calcium voltage-gated channel and G protein, respectively. These proteins interact with the nicotinic acetylcholine receptor, suggesting that methylation levels at these CpGs might be reactive to nicotine exposure. All 13 CpGs have been previously associated with smoking in unrelated individuals and data from monozygotic pairs discordant for former smoking indicated that methylation patterns are to a large extent reversible upon smoking cessation. We further showed that differences in smoking level exposure for monozygotic twins who are both current smokers but differ in the number of cigarettes they smoke are reflected in their DNA methylation profiles. In conclusion, by analysing data from monozygotic twins, we robustly demonstrate that DNA methylation level in human blood cells is reactive to cigarette smoking.

## Background

Epigenome-wide association studies (EWAS) have identified robust differences in DNA methylation between smokers and non-smokers[1]. In a meta-analysis of blood-based DNA methylation studies (N=15,907 individuals; the largest EWAS of smoking to date), 2,623 CpG sites passed the Bonferroni threshold for genome-wide significance in a comparison of current and never smokers[2]. Based on comparison with loci identified in large genome-wide association studies (GWAS), differentially methylated sites were significantly enriched in genes implicated in well-established smoking-associated diseases, such as cancer, cardiovascular disease, inflammatory disease and lung disease, as well as in genes associated with schizophrenia and educational attainment[2]. It has been hypothesized that smoking-induced methylation changes might also contribute to the addictive effect of smoking[3].

Importantly, smoking-associated DNA methylation levels, as established in human EWA studies, may reflect different mechanisms. They may reflect causal effects of smoking on methylation, causal effects of methylation on smoking behaviour, methylation differences associated with epiphenomena of other exposures that correlate with smoking (e.g. alcohol use[4]), or they may reflect a shared genetic predisposition to smoking and methylation level. To distinguish these different mechanisms requires incisive study designs[5]. Establishing whether methylation levels in smokers revert to levels of never smokers upon smoking cessation is a first step. A previous study of 2,648 former smokers with cross-sectional methylation data from the Framingham Heart Study suggested that methylation levels at most CpGs return to the level of never smokers within five years after quitting smoking, but 36 CpGs were still differentially methylated in former smokers, who had quit smoking for 30 years[2]. In the large EWAS meta-analysis of smoking[2], 185 CpGs were differentially methylated between former and never smokers (compared to 2623 between current and never smokers). In addition, differences between former and never smokers were smaller than between current and never smokers. Reversible DNA methylation patterns may suggest that DNA methylation is reactive to smoking. However, it is also possible that the different methylation level in current smokers reflects a higher genetic liability to smoking behavior (that makes them more likely to initiate and keep smoking). Similarly, differences between former smokers and never smokers could reflect that smoking has caused a persistent methylation change but can also be driven by genetic factors.

In population-based studies, smoking cases and non-smoking individuals may differ on many aspects, including their genetic predisposition to smoking. On the other hand, monozygotic twins are genetically identical (except for *de novo* mutations, but these are rare[6,7]), and are matched on sex, age and early environment. They have been exposed to similar prenatal conditions, which may include second hand smoke from smoking mothers and others. Therefore, smoking discordant monozygotic twin pairs offer a unique opportunity to assess smoking-reactive DNA methylation patterns[5,8]. Despite the large number of previous population-based smoking EWASs, only one previous study compared genome-wide DNA methylation in smoking discordant monozygotic twin pairs[9]. This study analysed whole blood Illumina 450k array methylation data from 20 discordant pairs, and reported 22 top loci, many of which had been previously associated with cigarette smoking in previous studies. However, following the correction for multiple testing, none of the differentially methylated loci were statistically significant, and this previous twin study did not examine reversibility of smoking effects, i.e., where methylation status changes again following smoking cessation.

Here, we analyse unique data from a large cohort of monozygotic twin pairs. This cohort is sufficiently large to include current smoking discordant and concordant pairs, as well as pairs discordant for former smoking. These groups allow identification of loci that are reactive to smoking, and examination of the extent to which effects are reversible upon quitting smoking. Monozygotic pairs in which both twins are current smokers, but who differ in quantity smoked, enable examination of the effects of smoking intensity. Finally, concordant pairs who never smoked allow assessment of the amount of DNA methylation variation at smoking-reactive loci that is due to non-genetic sources of variation other than smoking. In secondary enrichment analyses, we examined whether smoking-reactive methylation patterns are enriched 1) at loci detected in previous epigenome-wide association studies of other traits and exposures, 2) at loci detected in a previous large Genome-wide Association Study meta-analysis of smoking initiation[10] – these loci are presumed to have a causal effect on smoking behaviour, and 3) within Gene Ontology and Kegg Pathways. Finally, we examined the relationship between DNA methylation and RNA transcript levels in blood for smoking-reactive loci.

## Methods

### Participants

In the Netherlands Twin Register (NTR), DNA methylation data are available for 3087 whole blood samples from 3055 individuals in twin families, as described in detail previously[11]. The samples were obtained from twins and family members, who participated in NTR longitudinal survey studies[12] and the NTR biobank project[13]. In the current study, methylation data from monozygotic twin pairs were analysed. Among 768 monozygotic twin pairs with genome-wide methylation data and information on smoking and covariates, we identified the following discordant pairs: 53 discordant pairs, in which one twin was a current smoker at blood draw and the other never smoked, 72 discordant pairs, in which one twin was a former smoker at blood draw and the other never smoked, 66 discordant pairs of which one twin was a former smoker and the other a current smoker at blood draw. In addition, we identified the following concordant pairs: 83 twin pairs concordant for current smoking, 88 twin pairs concordant for former smoking, and 406 concordant twin pairs who never smoked. Informed consent was obtained from all participants. The study was approved by the Central Ethics Committee on Research Involving Human Subjects of the VU University Medical Centre, Amsterdam, an Institutional Review Board certified by the U.S. Office of Human Research Protections (IRB number IRB00002991 under Federal-wide Assurance- FWA00017598; IRB/institute codes, NTR 03-180).

### Peripheral blood DNA methylation and cell counts

Genome-wide DNA methylation in whole blood was measured by the Human Genomics facility (HugeF) of ErasmusMC, the Netherlands (http://www.glimdna.org/). DNA methylation was assessed with the Infinium HumanMethylation450 BeadChip Kit (Illumina, San Diego, CA, USA). Genomic DNA (500□ng) from whole blood was bisulfite treated using the Zymo EZ DNA Methylation kit (Zymo Research Corp, Irvine, CA, USA), and 4□μl of bisulfite-converted DNA was measured on the Illumina 450k array[14] following the manufacturer’s protocol. A custom pipeline for quality control and normalization of the methylation data was developed by the BIOS consortium. First, sample quality control was performed using MethylAi[15]. Next, probe filtering was applied with DNAmArray[16] to remove: ambiguously mapped probes[17], probes with a detection *P* value□>□0.01, or bead number < 3, or raw signal intensity of zero. After these probe filtering steps, probes and samples with a success rate□<□95% were removed. Next, the DNA methylation data were normalized using functional normalization[18], as implemented in DNAmArray[16] using the cohort-specific optimum number of control probe-based principal components. Probes containing a SNP, identified in a DNA sequencing project in the Dutch population[19], within the CpG site (at the C or G position) were excluded irrespective of minor allele frequency, and only autosomal probes were analysed, leading to a total number of 411,169 methylation sites. The following subtypes of white blood cells were counted in blood samples: neutrophils, lymphocytes, monocytes, eosinophils, and basophils[13].

### Smoking

Information on smoking behavior was obtained by interview during the home visit for blood collection as part of the NTR biobank project (2004–2008 and 2010-2011). Participants were asked: “Did you ever smoke?”, with answer categories: (1) no, I never smoked (2) I’m a former smoker (3) yes. Current smokers were asked how many years they smoked and how many cigarettes per day they smoked, while ex-smokers were asked how many years ago they quit, for how many years they smoked and how many cigarettes per day they smoked. Data were checked for consistencies and missing data were completed by linking this information to data from surveys filled out close to the time of biobanking within the longitudinal survey study of the NTR. Pack-years were calculated as the (number of cigarettes smoked per day)/20□×□number of years smoked.

### Statistical analyses

#### Overview and hypotheses

All analyses were performed in R[20]. Analyses were performed in six groups of monozygotic twin pairs (**Fig. 1**). To identify DNA methylation differences in smoking discordant monozygotic twin pairs, we first compared the twin pairs, in which one twin had never smoked, and the other was a current smoker at the time of blood sampling. Second, to identify which of these DNA methylation differences might be reversible, we analysed data from 1) monozygotic pairs in which one twin had never smoked, and the other was a former smoker at the time of blood sampling, 2) from monozygotic pairs in which one twin was a current smoker, and the other was a former smoker at the time of blood sampling, and 3) from monozygotic pairs who were both former smokers. Third, to quantify within-pair methylation differences that occur by chance alone, we compared the within-pair differences monozygotic twins concordant for never having smoked. Forth, data monozygotic twins concordant for current smoking were analysed to examine the effects of smoking intensity. Our hypotheses were as follows: 1) if DNA methylation level is reactive to cigarette smoking, methylation differences will be present between smokers and non-smokers after ruling out genetic differences, i.e. in smoking discordant monozygotic twins, and these differences will be larger than in monozygotic pairs concordant for never smoking, 2) if DNA methylation patterns are reversible upon quitting smoking, methylation differences (ΔM) in monozygotic pairs will show the following pattern: ΔM discordant current-never >ΔM discordant current-former and ΔM discordant former-never >ΔM concordant never, 3) a correlation between time since quitting smoking and ΔM in pairs discordant for former smoking is consistent with a gradual reversibility of methylation levels upon quitting smoking, 4) a correlation between ΔM and the difference in number of cigarettes smoked per day in smoking concordant pairs is consistent with smoking-reactive methylation patterns that show a dose-response relationship with amount of cigarettes smoked.

**Figure 1.**
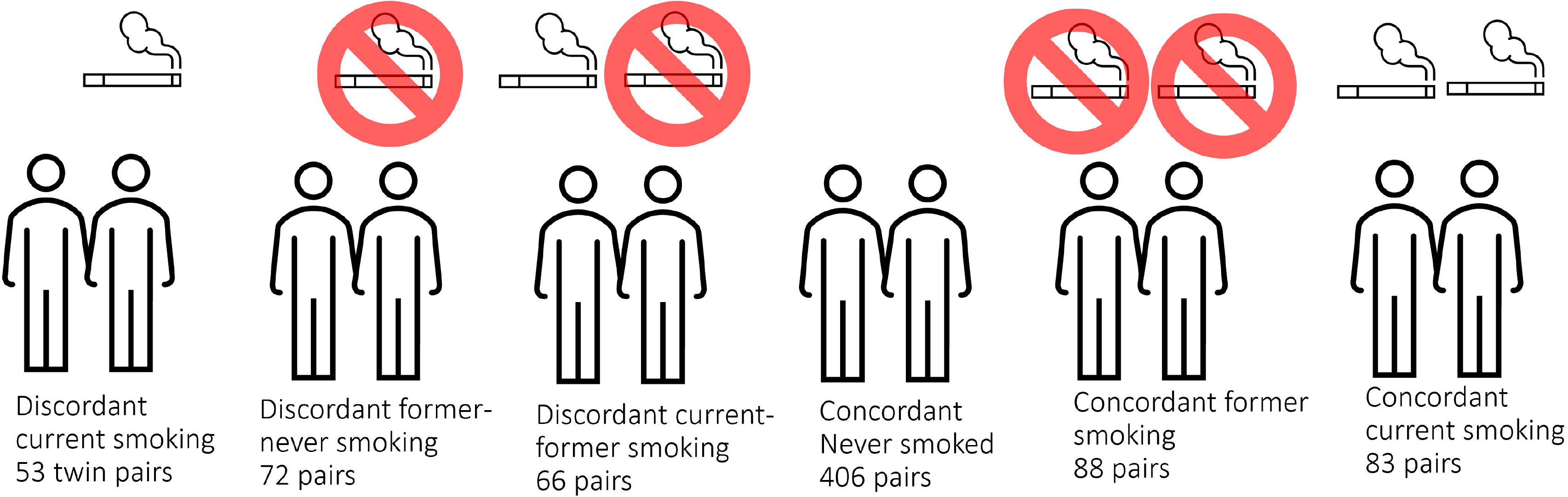
DNA methylation analysis in smoking discordant and smoking concordant monozygotic twin pairs. Blood DNA methylation profiles (Illumina 450k array) from six groups of monozygotic twin pairs were analysed.

#### Epigenome-wide association study

In the entire data set of 3087 blood samples, we used linear regression analysis to correct the DNA methylation levels (β -values) for HM450k array row, bisulphite plate (dummy-coding) and white blood cell percentages (% neutrophils, % monocytes, and % eosinophils). White blood cell percentages were included to account for variation in cellular composition between whole blood samples. Lymphocyte percentage was not included in models because it was strongly correlated with neutrophil percentage (r = −0.93), and basophil percentage was not included because it showed little variation between subjects, with a large number of subjects having 0% of basophils. We did not adjust for sex and age, because monozygotic twins have the same sex and age. The residuals from this regression analysis were used in the within-pair EWAS analyses. Specifically, the residuals were used as input for paired *t*-tests to compare the methylation of the smoking twins with that of their non-smoking cotwins. Similarly, paired *t*-tests were applied to data from smoking concordant pairs, and to test for differences in white blood cell profiles and smoking frequency. Statistical significance was assessed following Bonferroni correction for the number of methylation sites tested (α□=□00.05/411,169□=□1.2 x 10^-7^). A previous power analysis for DNA methylation studies in discordant monozygotic twins indicated that with 50 discordant pairs, there’s 80% power to detect methylation differences of 15% (at epigenome-wide significance; i.e. following multiple testing correction)[21]. Power quickly drops for smaller effect sizes; e.g. with 50 discordant pairs, the power to detect a 10% methylation difference is 10% and the power to detect a methylation difference of 5% approaches alpha[21].

#### Dose-response relationships

For significant CpGs from the EWAS of discordant monozygotic twin pairs, we examined dose-response relationships in smoking concordant pairs (both twins were current smokers) by correlating within-pair differences in DNA methylation with within-pair differences in smoking packyears and cigarettes per day. Secondly, in twin pairs discordant for former smoking (one twin never smoked and the other one is a former smoker), we correlated and plotted within-pair differences in DNA methylation with the time since quitting smoking to assess the relationship between time since quitting smoking and reversal of methylation differences within monozygotic twin pairs.

#### Enrichment analyses

We used the EWAS Toolkit from the EWAS atlas[22] to perform enrichment analyses of Gene Ontology Terms, Kegg Pathways, and previously associated traits among top-sites from the EWAS in discordant monozygotic twin pairs (current versus never). With the trait enrichment tool of the EWAS analysis, we tested for enrichment of all traits (680) that were present in the atlas on April 26, 2022. Because the software requires a minimum of 20 input CpGs, we selected the top 20 CpGs from the EWAS in discordant monozygotic pairs for the enrichment analyses using the EWAS toolkit.

To study overlap of EWAS signal with genetic findings for smoking, we considered the largest GWAS meta-analysis of smoking phenotypes, which is the meta-analysis of smoking initiation by the GWAS and Sequencing Consortium of Alcohol and Nicotine use (GSCAN)[10]. We obtained leave-one out meta-analysis results with NTR excluded. From the GWAS, we selected all SNPs with a p-value <5.0×10^-8^ and determined the distance of each Illumina 450k methylation site to each SNP. We then tested whether methylation sites within 1 Mb of genome-wide significant SNPs from the GWAS showed a stronger signal in the within-pair EWAS of smoking discordant monozygotic pairs compared to other genome-wide methylation sites, by regressing the EWAS test statistics on a variable (“GWAS locus”) indicating if the CpG is located within a 1 Mb window from SNPs associated with smoking initiation (1□=□yes, 0□=□no):

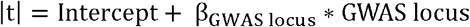

Where |t| represents the absolute t-statistic from the paired t-test comparing within-pair methylation differences in smoking discordant pairs and β_GWAS locus_ represents the estimate for GWASlocus, i.e. the change in the t-test statistic associated with a one-unit change in the variable GWAS locus (e.g. being within 1 Mb of SNPs associated with smoking initiation). For each enrichment test, bootstrap standard erors were computed with 2000 bootstraps with the R-package ‘simpleboot’.

### Gene Expression

For significant CpGs from the EWAS of discordant monozygotic twin pairs (current versus never), we examined whether the DNA methylation was associated with gene expression levels in *cis*. To this end, we used an independent whole blood RNA-sequencing dataset from the Biobank-based Integrative Omics Study (BIOS) consortium that did not include NTR, and tested associations between genome-wide CpGs and transcripts in cis (<250 kb), as described in detail previously[23]. In short, methylation and expression levels in whole-blood samples (n=2,101) were quantified with Illumina Infinium HumanMethylation450 BeadChip arrays and with RNA-seq (2×50bp paired-end, Hiseq2000, >15M read pairs per sample). For each target CpG (epigenome-wide significant DMPs), we identified transcripts in cis (<250 kb), for which methylation levels were significantly associated with gene expression levels at the experiment-wide threshold applied by this study (FDR<5.0%), after regressing out mQTL and eQTL effects. We also examined whether significant CpGs from the EWAS of discordant monozygotic twin pairs mapped to genes that were previously reported to be differentially expressed in monozygotic pairs of which one twin never smoked, and the other was a current smoker at the time of blood sampling (based on Affymetrix U219 array data; n□=□56 pairs; note: the 53 discordant pairs included in the current study of DNA methylation are a sub-set of the 56 discordant pairs included in the study of gene expression)[5].

## Results

Descriptives of the smoking discordant and concordant monozygotic twin pairs are given in **table 1**. In twin pairs discordant for current smoking status (N=53 pairs, mean age=33 years), the smoking twin on average smoked 8.9 cigarettes per day at the time of blood sampling, and had an average smoking history equivalent to 6.8 pack years. The epigenome-wide association study (EWAS) analysis in pairs discordant for current smoking status identified 13 epigenome-wide significant (P<1.20×10^-7^) differentially methylated positions (DMPs; **Fig. 2a**). Genome-wide test statistics were not inflated (**Additional file 1**). Absolute differences in methylation ranged from 2.5-13% (0.025-0.13 on the methylation ß- value scale), with a mean of 5.4% (**Table 2**). Eight of the 13 CpGs (61.5%) showed lower methylation in the current smoking twins compared to their non-smoking twins.

**Figure 2.**
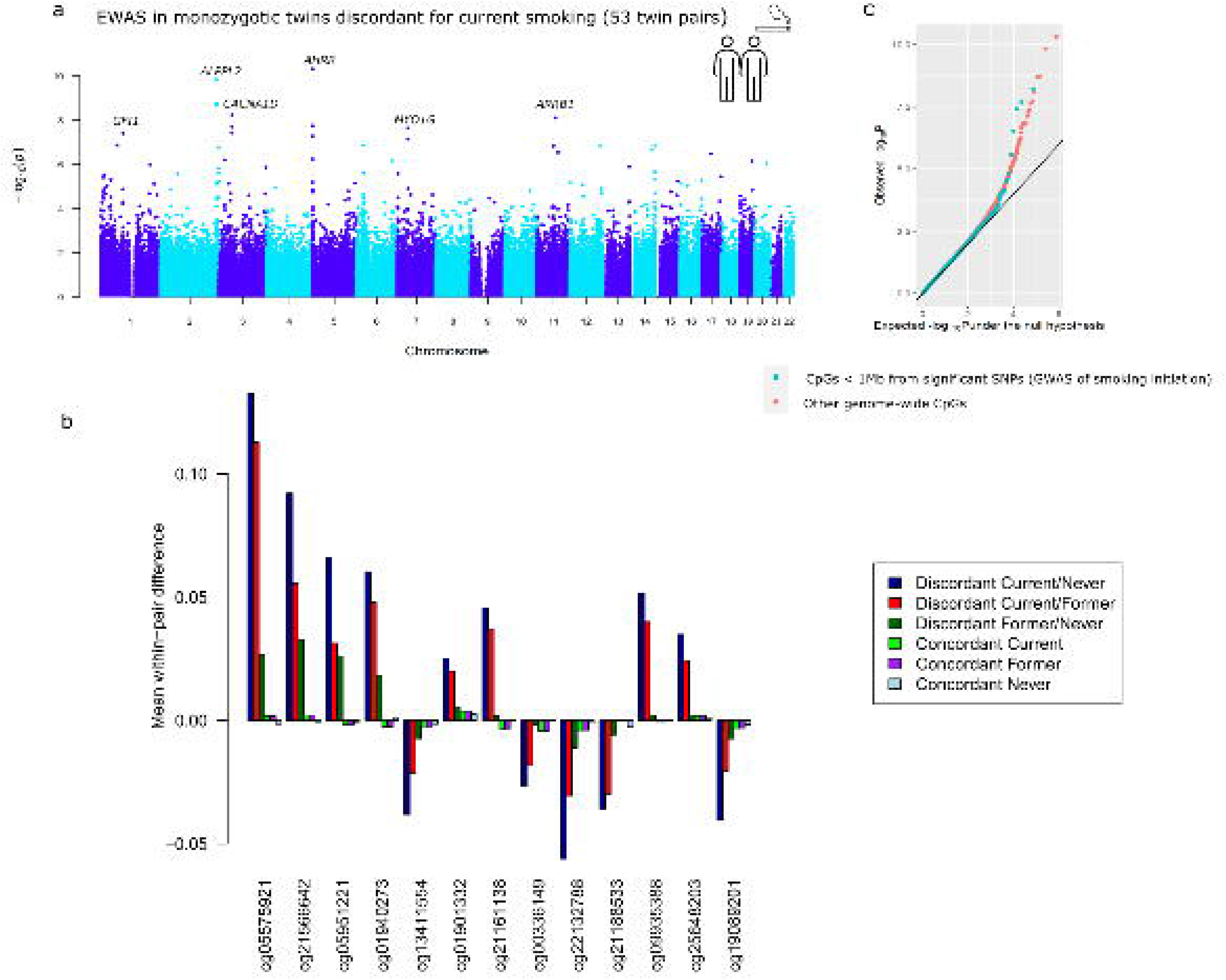
Top differentially methylated loci identified in monozygotic twin pairs discordant for current smoking. a) Manhattan plot of the EWAS in 53 smoking discordant monozygotic twin pairs (current versus never). The red horizontal line denotes the epigenome-wide significance threshold (Bonferroni correction) and 13 CpGs with significant differences are highlighted. b) Mean within-pair differences in monozygotic twin pairs at the 13 CpGs that were epigenome-wide significant in smoking discordant monozygotic pairs. Mean within-pair differences of the residuals obtained after correction of methylation beta-values for covariates are shown for 53 monozygotic pairs discordant for current/never smoking, 66 monozygotic pairs discordant for current/former smoking, 72 monozygotic pairs discordant for former/never smoking, 83 concordant current smoking monozygotic pairs, 88 concordant former smoking monozygotic pairs, and 406 concordant never smoking monozygotic pairs. c) QQ-plot showing p-values from the EWAS in 53 smoking discordant monozygotic twin pairs (current versus never). P-values for CpGs located nearby significant SNPs from the GWAS of smoking initiation are plotted in blue and all other genome-wide CpGs are plotted in orange.

**Table 1.** Descriptive statistics for smoking discordant and concordant monozygotic twin pairs

|  | Discordant Current/never<br>(53 pairs) |  |  |  | Discordant Former/never<br>(72 pairs) |  |  |  | Discordant Current/former<br>(66 pairs) |  |  |  | Concordant Current<br>(83 pairs) |  |  |  | Concordant never<br>(406 pairs) |  |  |  | Concordant former<br>(88 pairs) |  |  |  |
| --- | --- | --- | --- | --- | --- | --- | --- | --- | --- | --- | --- | --- | --- | --- | --- | --- | --- | --- | --- | --- | --- | --- | --- | --- |
|  | Current<br>smoker | Never-<br>smoker | Mean<br>diff | P-value | Former<br>smoker | Never-<br>smoker | Mean<br>diff | P-value | Current<br>smoker | Former<br>smoker | Mean<br>diff | P-value | Twin 1 | Twin<br>2 | Mean<br>diff | P-value | Twin 1 | Twin 2 | Mean<br>diff | P-value | Twin 1 | Twin 2 | Mean diff | P-value |
| % Female pairs | 60.4% | 60.4% | n.a. | n.a. | 77.80% | 77.80% | n.a. | n.a. | 69.7% | 69.7% | n.a. | n.a. | 61.4% | 61.4% | n.a. | n.a. | 73.6% | 73.6% | n.a. | n.a. | 64.8% | 64.8% | n.a. | n.a. |
| age at blood sampling, mean<br>(SD) | 33.1<br>(8.0) | 33.0<br>(7.9) | 0.10 | 0.34 | 41.4<br>(13.2) | 41.4<br>(13.1) | 0.02 | 0.83 | 42.2<br>(12.6) | 42.2<br>(12.5) | -0.06 | 0.45 | 33.8<br>(10.3) | 33.9<br>(10.5) | -0.12 | 0.10 | 33.1<br>(11.3) | 33.0<br>(11.2) | 0.06 | 0.08 | 45.2<br>(13.4) | 45.2<br>(13.4) | 0.09 | 0.29 |
| Cigarettes per day at blood<br>sampling, mean (SD), N<br>missings | 8.9<br>(6.4), 6 | n.a. | n.a. | n.a. | n.a. | n.a. | n.a. | n.a. | 11.9<br>(7.2), 9 | n.a. | n.a. | n.a. | 11.1<br>(7.0), 2 | 10.9<br>(6.9),<br>1 | 0.00 | 1.00 | n.a. | n.a. | n.a. | n.a. | n.a. | n.a. | n.a. | n.a. |
| Packyears, mean (SD), N<br>missings | 6.8<br>(7.0),<br>13 | n.a. | n.a. | n.a. | 5.9<br>(11.1),<br>15 | n.a. | n.a. | n.a. | 13.6<br>(13.2), 9 | 9.3<br>(8.7), 10 | 4.2 | 0.02 | 9.7<br>(9.3), 10 | 8.3<br>(7.6),<br>9 | 0.22 | 0.82 | n.a. | n.a. | n.a. | n.a. | 10.6<br>(11.5), 7 | 9.8<br>(10.4), 1<br>1 | 0.78 | 0.55 |
| Years since quitting smoking,<br>mean (SD), N missings | n.a. | n.a. | n.a. | n.a. | 13.5<br>(11.4),<br>9 | n.a. | n.a. | n.a. | n.a. | 9.0<br>(10.2), 2 | n.a. | n.a. | n.a. | n.a. | n.a. | n.a. | n.a. | n.a. | n.a. | n.a. | 11.9<br>(9.1), 8 | 13.6<br>(11.8), 7 | -1.62 | 0.20 |
| Percentage monocytes, mean<br>(SD), N missings | 8.0<br>(2.3), 0 | 8.5<br>(2.4), 0 | -0.44 | 0.19 | 8.6<br>(2.0), 0 | 8.3<br>(1.8), 0 | 0.29 | 0.19 | 8.6<br>(1.9), 0 | 9.2<br>(3.1), 0 | -0.57 | 0.14 | 8.3<br>(2.1), 0 | 8.1<br>(1.9),<br>0 | 0.20 | 0.36 | 8.5<br>(2.0), 0 | 8.5<br>(2.2), 0 | 0.03 | 0.75 | 8.4<br>(2.4), 0 | 8.5<br>(2.4), 0 | -0.06 | 0.75 |
| Percentage lymphocytes, mean<br>(SD), N missings | 35.0<br>(8.9), 0 | 35.9<br>(10.0) | -0.94 | 0.50 | 33.6<br>(8.5), 0 | 34.0<br>(8.7) | -0.37 | 0.77 | 35.8<br>(8.2), 0 | 35.6<br>(8.5), 0 | 0.25 | 0.84 | 33.7<br>(8.3), 0 | 34.1<br>(8.3),<br>0 | -0.44 | 0.67 | 36.3<br>(8.4), 0 | 36.2<br>(8.4), 0 | 0.04 | 0.92 | 35.0<br>(7.7), 0 | 34.1<br>(8.4), 0 | 0.95 | 0.26 |
| Percentage neutrophils, mean<br>(SD), N missings | 53.4<br>(9.5), 0 | 52.1<br>(9.8), 0 | 1.34 | 0.38 | 54.4<br>(9.1), 0 | 54.5<br>(9.1), 0 | -0.08 | 0.95 | 51.6<br>(9.0), 0 | 51.7<br>(8.9), 0 | 0-0.06 | 0.96 | 53.7<br>(8.9), 0 | 53.7<br>(9.3),<br>0 | 0.03 | 0.98 | 51.8,<br>(8.7), 0 | 51.9<br>(9.3), 0 | -0.08 | 0.86 | 52.8<br>(8.2), 0 | 53.7<br>(8.4), 0 | -0.84 | 0.35 |
| Percentage eosinophils, mean<br>(SD), N missings | 3.1<br>(2.5), 0 | 3.1<br>(2.1), 0 | .05 | 0.91 | 3.1<br>(1.9), 0 | 2.9<br>(2.0), 0 | 0.15 | 0.53 | 3.3<br>(1.9), 0 | 3.1<br>(1.7), 0 | 0.21 | 0.33 | 3.4<br>(2.2), 0 | 3.4<br>(1.8),<br>0 | -0.03 | 0.91 | 2.9<br>(1.8), 0 | 2.9<br>(1.9), 0 | -0.04 | 0.66 | 3.1<br>(1.9), 0 | 3.2<br>(2.4), 0 | -0.06 | 0.77 |
| Percentage basophils, mean<br>(SD), N missings | 0.5<br>(0.7), 0 | 0.5<br>(0.7), 0 | -0.01 | 0.96 | 0.3<br>(0.3), 0 | 0.4<br>(0.5), 0 | -0.02 | 0.76 | 0.6<br>(0.9), 0 | 0.4<br>(0.4), 0 | 0.18 | 0.12 | 0.9<br>(3.1), 0 | 0.6<br>(1.1),<br>0 | 0.25 | 0.49 | 0.5<br>(0.7), 0 | 0.4<br>(0.7), 0 | 0.06 | 0.21 | 0.6<br>(1.1), 0 | 0.6<br>(0.9), 0 | -0.01 | 0.97 |

**Table 2.**
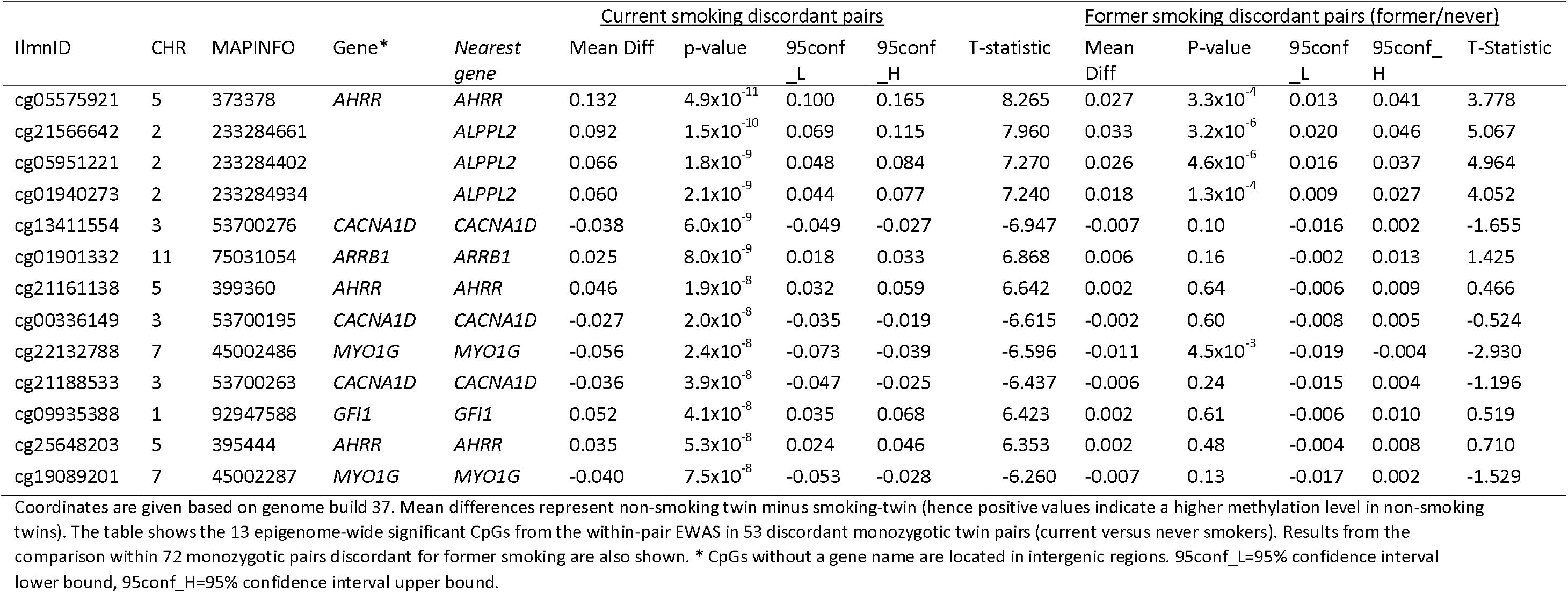
Epigenome-wide significant differentially methylated CpGs in monozygotic pairs discordant for current smoking status

In twin pairs discordant for former smoking (N=72 pairs, mean age=41 years), the twins, who used to smoke, had quit smoking on average 14 years ago, while the other twins had never initiated regular smoking. In this group, no epigenome-wide significant DMPs were identified, and within-pair differences at the 13 significant DMPs identified in the previous analysis were diminished (average reduction: 81%, range=61-96%; **Fig. 2b, Table 2**). By contrast, in twin pairs of which one twin was a current smoker at blood draw and the co-twin had quit smoking (on average 9 years, ago, N=66 pairs), the reduction of within-pair differences at the 13 top CpGs was much smaller (on average, 31%, rage 15-52%; **Fig. 2b, Additional file 2**), and 5 of the 13 DMPs identified by comparing current and never smoking twins were also epigenome-wide significant in this group. Furthermore, 5 additional epigenome-wide CpGs were identified in current/former smoking discordant pairs (**Additional file 3**). **Fig. 2b** illustrates the pattern of within-pair differences at the 13 top DMPs identified in current/never discordant monozygotic pairs: largest differences in current/never smoking discordant pairs, smaller differences in former/never discordant pair, and current/former discordant pairs are intermediate. Differences are smallest within smoking concordant pairs. This pattern is in line with smoking-associated methylation patterns in blood cells being to a large extent reversible upon quitting smoking.

Distributions of within-pair differences in smoking-discordant and concordant pairs for the top 1000 CpGs of the EWAS in discordant pairs are shown in **Fig. 3a**. The distributions illustrate that differences are largest, as expected, within monozygotic twin pairs discordant for current smoking (current/never smoking pairs), followed by discordant current/former smoking discordant pairs, followed by former/never smoking discordant monozygotic twin pairs. Monozygotic pairs concordant for current smoking also show notable within-pair differences at these CpGs that are substantially larger compared to monozygotic pairs concordant for never smoking (**Fig. 3a**). This could be explained by within-pair differences in the number of cigarettes smoked by monozygotic twins who were concordant for current smoking. The twin correlations in current smoking monozygotic twin pairs were *r*=0.50, p= 2.2×10^-6^ for cigarettes per day (**Fig. 3b**) and *r*=0.43, p=3.2 x10^-4^ for packyears, respectively. Within-pair differences in DNA methylation at the 13 top-CpGs correlated with within-pair differences in the number of cigarettes smoked per day (mean absolute *r=* 0.38, range (for different CpGs): −0.56-0.41; **Table 3**, **Fig. 3c**) and with within-pair differences in packyears (mean absolute *r*= 0.46; range: −0.65-0.42; **Table 3**). In twin pairs discordant for former smoking, within-pair differences in DNA methylation at the 13 top-CpGs were weakly correlated with time since quitting smoking (mean *r*= −0.11, range=-0.28- 0.05, **Additional file 4**). Based on scatterplots of the within-pair methylation differences against time since quitting smoking (**Fig. 3d**), we hypothesized that the lack of a strong correlation with time since quitting smoking might be explained by most of the reversal taking place within the first years after quitting smoking. We therefore repeated the analysis restricting to those pairs of which the smoking twin had quit smoking less than 5 years ago (N=15 pairs). In this group, within-pair differences in DNA methylation at the 13 top-CpGs were on average more strongly correlated with time since quitting smoking (mean *r*= −0.16, range=-0.48- 0.23) but the sample size was greatly reduced and correlations were non-significant.

**Figure 3.**
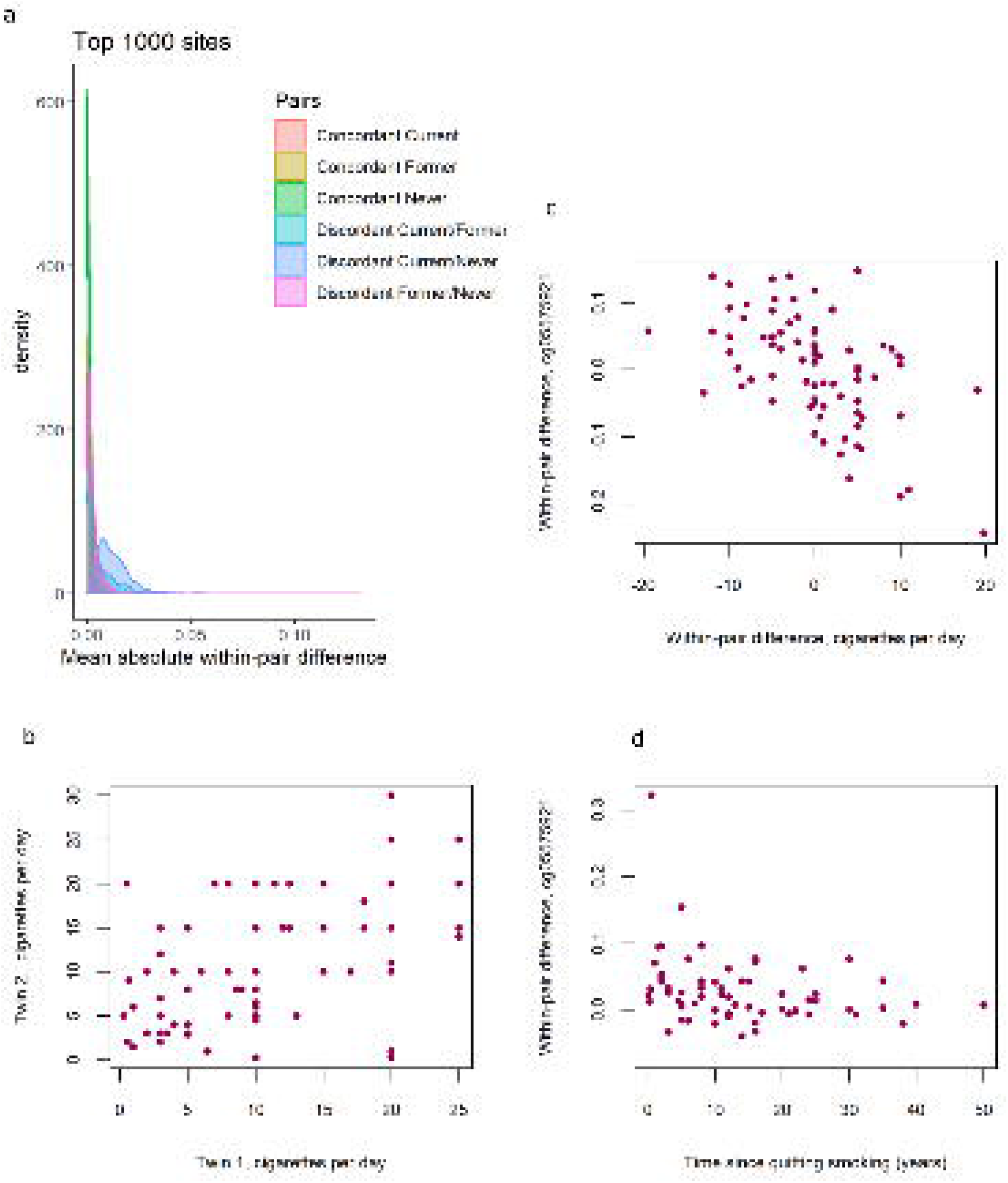
DNA methylation differences in smoking discordant and smoking concordant pairs. a) Distributions of the mean absolute within-pair differences in discordant and concordant pairs at the top 1000 CpGs with the lowest p-value from the EWAS in discordant monozygotic pairs (current versus never smokers). b) Scatterplot of cigarettes smoked per day in 80 concordant current smoking monozygotic pairs with complete data c) Scatterplot of within-pair differences in cigarettes smoked per day versus DNA methylation at cg05575921 (*AHRR*) in 80 concordant current smoking monozygotic pairs with complete data. d) Scatterplot of within-pair differences in DNA methylation at cg05575921 (*AHRR*) versus time since quitting smoking (years) in 63 pairs discordant for former smoking.

**Table 3.** Correlations of within-pair differences in DNA methylation with within-pair differences in cigarettes per day and packyears in 83 concordant current smoking monozygotic pairs

| cgid | CHR | Position | Gene | Nearest gene | cigarettes per day |  | Packyears |  |
| --- | --- | --- | --- | --- | --- | --- | --- | --- |
|  |  |  |  |  | *r* | p-value | *r* | p-value |
| cg05575921 | 5 | 373378 | AHRR | AHRR | -0.52 | 5.9x10<sup>-7</sup> | -0.55 | 1.2x10<sup>-6</sup> |
| cg21566642 | 2 | 233284661 |  | ALPPL2 | -0.49 | 4.7x10<sup>-6</sup> | -0.56 | 8.1x10<sup>-7</sup> |
| cg05951221 | 2 | 233284402 |  | ALPPL2 | -0.44 | 4.7x10<sup>-5</sup> | -0.56 | 9.0x10<sup>-7</sup> |
| cg01940273 | 2 | 233284934 |  | ALPPL2 | -0.56 | 7.0x10<sup>-8</sup> | -0.65 | 2.0x10<sup>-9</sup> |
| cg13411554 | 3 | 53700276 | CACNA1D | CACNA1D | 0.27 | 1.4x10<sup>-2</sup> | 0.32 | 7.4x10<sup>-3</sup> |
| cg01901332 | 11 | 75031054 | ARRB1 | ARRB1 | -0.23 | 3.8x10<sup>-2</sup> | -0.34 | 5.5x10<sup>-3</sup> |
| cg21161138 | 5 | 399360 | AHRR | AHRR | -0.52 | 8.4x10<sup>-7</sup> | -0.52 | 7.4x10<sup>-6</sup> |
| cg00336149 | 3 | 53700195 | CACNA1D | CACNA1D | 0.36 | 1.0x10<sup>-3</sup> | 0.42 | 3.5x10<sup>-4</sup> |
| cg22132788 | 7 | 45002486 | MYO1G | MYO1G | 0.41 | 2.1x10<sup>-4</sup> | 0.41 | 8.4x10<sup>-4</sup> |
| cg21188533 | 3 | 53700263 | CACNA1D | CACNA1D | 0.32 | 4.1x10<sup>-3</sup> | 0.38 | 1.4x10<sup>-3</sup> |
| cg09935388 | 1 | 92947588 | GFI1 | GFI1 | -0.42 | 9.1x10<sup>-5</sup> | -0.59 | 1.4x10<sup>-7</sup> |
| cg25648203 | 5 | 395444 | AHRR | AHRR | -0.28 | 1.2x10<sup>-2</sup> | -0.42 | 4.3x10<sup>-4</sup> |
| cg19089201 | 7 | 45002287 | MYO1G | MYO1G | 0.15 | 1.8x10<sup>-1</sup> | 0.27 | 2.6x10<sup>-2</sup> |

All 13 differentially methylated CpGs identified in current smoking-discordant pairs have been previously associated with smoking. To the study the overlap of methylation differences between smoking discordant twins with loci that have a causal effect on smoking, we considered the largest GWAS meta-analysis of smoking phenotypes, the meta-analysis of smoking initiation by the GWAS and Sequencing Consortium of Alcohol and Nicotine use (GSCAN)[10]. Three of the 13 epigenome-wide significant DMPs detected in smoking-discordant monozygotic pairs (cg13411554, cg00336149, and cg21188533 in *CACNA1D*) are located within 1Mb of a GWAS locus associated with smoking initiation. The methylation sites within 1 Mb of genome-wide significant SNPs from the GWAS overall did not show a stronger signal in the within-pair EWAS of smoking discordant monozygotic pairs compared to other genome-wide methylation sites (beta=-0.002, se=0.004, p=0. 0.56, **Fig. 2c**).

We tested for enrichment of methylation sites previously associated with 680 traits reported in the EWAS atlas[22], among the top differentially methylated loci in smoking discordant pairs, which showed strong enrichment of smoking-related traits (**Additional file 5**). Enrichment analysis based on Kegg Pathways showed one significantly enriched pathway; Dopaminergic Synapse (hsa04728; **Additional file 6**), with 3 of the top differentially methylated loci in smoking discordant monozygotic pairs mapping to this pathway; *CACNA1D, GNG12*, and *ARRB1*. No significant enrichment was seen in GO pathways after multiple testing correction (**Additional file 7**).

To examine potential functional consequences of top DMPs, we used previously published data on whole-blood DNA methylation and RNA sequencing (n = 2,101 samples). At four of the 13 CpGs, DNA methylation level in blood was associated with the expression level of nearby genes (**Table 4**). At three CpGs, a higher methylation level correlated with lower expression level. None of the 13 CpGs overlapped with six genes that were differentially expressed in monozygotic pairs discordant for current smoking[5].

**Table 4.**
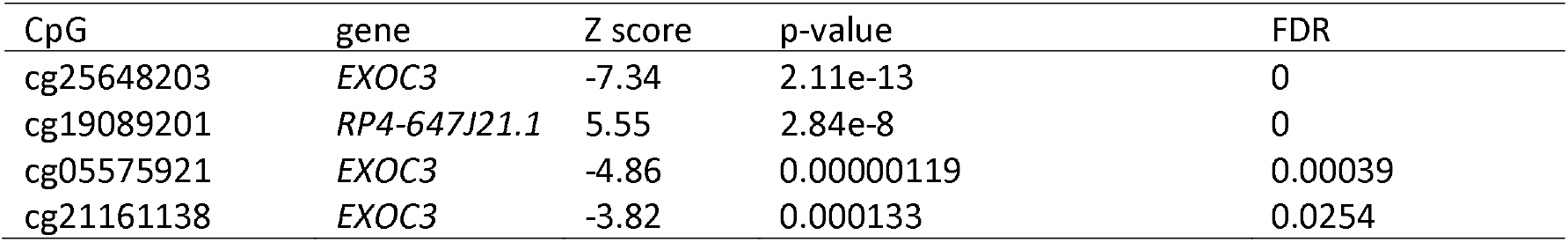
Significantly associated transcripts in cis for CpGs that are differentially methylated in smoking discordant monozygotic twin pairs

## Discussion

Previous epigenome-wide association studies (EWAS) have identified robust differences in DNA methylation between smokers and non-smokers at a number of loci. These differences may reflect true smoking-reactive DNA methylation patterns, but can also be driven by (genetic) confounding or reverse causation. We exploited a strong within-family design, i.e., the discordant monozygotic twin design[24], to identify smoking-reactive loci. By analysing whole blood genome-wide DNA methylation patterns in 53 monozygotic pairs discordant for current smoking, we found 13 CpGs with a difference in methylation level between the current smoking twin and the twin who never smoked. All 13 CpGs have been previously associated with smoking in unrelated individuals and in line with previous studies that compared unrelated smokers and controls[2], our data from monozygotic pairs discordant for former smoking also indicate that methylation patterns are to a large extent reversible upon smoking cessation. We further showed that differences in smoking level exposure for monozygotic twins who are both current smokers but differ in the number of cigarettes they smoke are reflected in their DNA methylation profiles.

The strongest smoking-associated loci typically detected in human blood EWAS are genes involved in detoxification pathways of aromatic hydrocarbons, such as *AHRR* and *CYP1A1[1]*, of which *AHRR* was also present among the top differentially methylated loci in our analysis of discordant twins. Mainstream tobacco smoke is a mixture of thousands of chemicals[25]. Although the effects of many of the compounds present in cigarette smoke are unknown, several mechanisms have been described through which cigarette smoking may affect global or gene-specific DNA methylation levels. These include DNA damage induced by certain compounds such as arsenic, chromium, formaldehyde, polycyclic aromatic hydrocarbons, and nitrosamines that all cause double-stranded breaks[26] (which causes increased methylation near repaired DNA) [27],[28], hypoxia induced by carbon monoxide[29] (causing global CpG island demethylation by disrupting methyl donor availability), and modulation of the expression level or activity of DNA-binding proteins, such as transcription factors[30]. Nicotine, presumed to be the major addictive compound in cigarette smoke (although other putative addictive compounds have also been described[31]), has gene regulatory effects. Binding of nicotine to nicotinic acetylcholine receptors causes downstream activation of cAMP response element binding protein, which is a key transcription factor for many genes[32]. In mouse brain, nicotine downregulates the DNA methyl transferase gene *Dnmt[33]*.

Importantly, effects of smoking on DNA methylation in brain cells have been hypothesized to contribute to addiction[3], but it is largely unknown to what extent addiction-related DNA methylation dynamics are captured in other tissues such as blood. Nicotinic receptors are especially abundant in the central and peripheral nervous system, but are also present in other tissues. In peripheral blood, nicotinic receptors are present on lymphocytes and polymorphonuclear cells[34], suggesting that EWA studies performed on blood cells might capture nicotine-reactive methylation patterns. Interesting in this regard is our finding that among the top differentially methylated CpGs in smoking discordant pairs are multiple CpGs in *CACNA1D and GNG12*, which encode subunits of a calcium voltage-gated channel and G protein, respectively; proteins that interact with the nicotinic acetylcholine receptor, and the related enrichment of Kegg Pathway dopaminergic neuron. Methylation levels at these CpGs might be reactive to nicotine exposure. Furthermore, the CpGs in *CACNA1D* are in proximity of a GWAS locus for smoking initiation, suggesting that this might be a locus that is not only reactive to smoking exposure, but may also contribute to smoking behaviour. Although it remains to be established if the epigenetic and genetic variation at this locus are functionally connected (i.e. have the same downstream consequences on gene expression), these results suggest that these CpGs can be interesting candidates for further studies into peripheral biomarkers of smoking addiction. Since we applied a discordant monozygotic twin design, the methylation differences identified at this locus in our study cannot be driven by mQTL effects of the SNPs associated with smoking.

The main strength of our study is the use of the discordant monozygotic twin design to examine the effects of smoking, because it rules out genetic confounding, as well as many other confounding factors. The value of studying smoking effects against an identical genetic background is clear if one considers that one of the most strongly associated genetic variants for nicotine dependence is located in the DNA methyltransferase gene *DNMT3B[35]*. This strongly implies a role for DNA methylation in nicotine addiction, but it also suggests that horizontal genetic pleiotropy might contribute to associations between DNA methylation and smoking in ordinary case-control EWASs, where differences in DNA methylation between unrelated smokers and non-smokers may reflect differences in genotype. Our analysis had adequate power to detect large effects (i.e., the top hits identified in typical smoking EWAS)[21]. Larger sample sizes are required to achieve adequate power to detect smaller effects. While the pattern of within-pair differences in current/never, current/former and former/never discordant monozygotic twin pairs was clearly in line with reversal of methylation patterns following smoking cessation, we did not find a strong correlation between within-pair differences in DNA methylation and time since quitting smoking in former smoking discordant pairs. If most reversal takes place gradually in the first view years after smoking cessation, it might require larger sample sizes of twin pairs discordant for recently quitting smoking to detect such a correlation. Larger samples sizes may be achieved by combining data from multiple twin cohorts in a meta-analysis. Common limitations that apply to many EWA studies including ours are that we only analysed DNA methylation data from blood and that the technique used to measure DNA methylation only covers a small sub-set of all CpG sites in the genome.

## Conclusion

In conclusion, we studied reactiveness of DNA methylation in blood cells to smoking and reversibility of methylation patterns upon quitting smoking in monozygotic twins. Analyses in special groups such as monozygotic twins are valuable to validate results from large population-based EWAS meta-analyses, or to train more accurate methylation scores for environmental exposures that are not confounded by genetic effects. Our results illustrate the potential to utilize DNA methylation profiles of monozygotic twins as a read out of discordant exposures at present and in the past.

## Availability of data and materials

The HumanMethylation450 BeadChip data from the NTR are available as part of the Biobank-based Integrative Omics Studies (BIOS) Consortium in the European Genome- phenome Archive (EGA), under the accession code EGAD00010000887. The pipeline for DNA methylation-array analysis developed by the Biobank-based Integrative Omics Study (BIOS) consortium is available here: https://molepi.github.io/DNAmArray_workflow/ (https://doi.org/10.5281/zenodo.3355292). All other analysis code is available upon request from the corresponding author.

## Acknowledgments

NTR warmly thanks all participants. We thank Conor Dolan for providing feedback on the manuscript. We acknowledge the contributions of the investigators of the BIOS consortium (Additional file 8): Bastiaan T. Heijmans, Peter A.C. ’t Hoen, Joyce van Meurs, Aaron Isaacs, Rick Jansen, Lude Franke, René Pool, Jouke J. Hottenga, Marleen MJ van Greevenbroek, Coen D.A. Stehouwer, Carla J.H. van der Kallen, Casper G. Schalkwijk, Cisca Wijmenga, Sasha Zhernakova, Ettje F. Tigchelaar, P. Eline Slagboom, Marian Beekman, Joris Deelen, Diana van Heemst, Jan H. Veldink, Leonard H. van den Berg, Cornelia M. van Duijn^4^, Bert A. Hofman^11^, Aaron Isaacs, André G. Uitterlinden, P. Mila Jhamai, Michael Verbiest, H. Eka D. Suchiman, Marijn Verkerk, Ruud van der Breggen, Jeroen van Rooij, Nico Lakenberg, Hailiang Mei, Maarten van Iterson, Michiel van Galen, Jan Bot, Dasha V. Zhernakova, Rick Jansen, Peter van ’t Hof, Patrick Deelen, Irene Nooren, Matthijs Moed, Martijn Vermaat, René Luijk, Marc Jan Bonder, Freerk van Dijk, Michiel van Galen, Wibowo Arindrarto, Szymon M. Kielbasa, Morris A. Swertz, Erik. W van Zwet.

## Author contributions

Phenotype acquisition or harmonization (including design/funding of contributing studies): GW, ECDG, DIB, MCN. Methylation data acquisition and/or QC: JVD, GW, DIB. Statistical Analysis: JVD. Interpretation of results: JVD, MCN, Draft manuscript preparation: JVD. All authors critically read and approved the final version of the manuscript.

