## Additional file 8 for "Genome-wide DNA methylation profiles in smoking discordant and concordant monozygotic twin pairs"

BIOS consortium members

**Management Team** Bastiaan T. Heijmans (chair)^1^, Peter A.C. ’t Hoen^2^, Joyce van Meurs^3^, Aaron Isaacs^4^, Rick Jansen^5^, Lude Franke^6^.

**Cohort collection** Dorret I. Boomsma^7^, René Pool^7^, Jenny van Dongen^7^, Jouke J. Hottenga^7^ (Netherlands Twin Register); Marleen MJ van Greevenbroek^8^, Coen D.A. Stehouwer^8^, Carla J.H. van der Kallen^8^, Casper G. Schalkwijk^8^ (Cohort study on Diabetes and Atherosclerosis Maastricht); Cisca Wijmenga^6^, Lude Franke^6^, Sasha Zhernakova^6^, Ettje F. Tigchelaar^6^ (LifeLines Deep); P. Eline Slagboom^1^, Marian Beekman^1^, Joris Deelen^1^, Diana van Heemst^9^ (Leiden Longevity Study); Jan H. Veldink^10^, Leonard H. van den Berg^10^ (Prospective ALS Study Netherlands); Cornelia M. van Duijn^4^, Bert A. Hofman^11^, Aaron Isaacs^4^, André G. Uitterlinden^3^ (Rotterdam Study).

**Data Generation** Joyce van Meurs (Chair)^3^, P. Mila Jhamai^3^, Michael Verbiest^3^, H. Eka D. Suchiman^1^, Marijn Verkerk^3^, Ruud van der Breggen^1^, Jeroen van Rooij^3^, Nico Lakenberg^1^.

**Data management and computational infrastructure** Hailiang Mei (Chair)^12^, Maarten van Iterson^1^, Michiel van Galen^2^, Jan Bot^13^, Dasha V. Zhernakova^6^, Rick Jansen^5^, Peter van ’t Hof^12^, Patrick Deelen^6^, Irene Nooren^13^, Peter A.C. ’t Hoen^2^, Bastiaan T. Heijmans^1^, Matthijs Moed^1^.

**Data Analysis Group** Lude Franke (Co-Chair)^6^, Martijn Vermaat^2^, Dasha V. Zhernakova^6^, René Luijk^1^, Marc Jan Bonder^6^, Maarten van Iterson^1^, Patrick Deelen^6^, Freerk van Dijk^14^, Michiel van Galen^2^, Wibowo Arindrarto^12^, Szymon M. Kielbasa^15^, Morris A. Swertz^14^, Erik. W van Zwet^15^, Rick Jansen^5^, Peter-Bram ’t Hoen (Co-Chair)^2^, Bastiaan T. Heijmans (Co-Chair)^1^.

1. Molecular Epidemiology Section, Department of Medical Statistics and Bioinformatics, Leiden University Medical Center, Leiden, The Netherlands

2. Department of Human Genetics, Leiden University Medical Center, Leiden, The Netherlands

3. Department of Internal Medicine, ErasmusMC, Rotterdam, The Netherlands

4. Department of Genetic Epidemiology, ErasmusMC, Rotterdam, The Netherlands

5. Department of Psychiatry, VU University Medical Center, Neuroscience Campus Amsterdam, Amsterdam, The Netherlands

6. Department of Genetics, University of Groningen, University Medical Centre Groningen, Groningen, The Netherlands

7. Department of Biological Psychology, VU University Amsterdam, Neuroscience Campus Amsterdam, Amsterdam, The Netherlands

8. Department of Internal Medicine and School for Cardiovascular Diseases (CARIM), Maastricht University Medical Center, Maastricht, The Netherlands

9. Department of Gerontology and Geriatrics, Leiden University Medical Center, Leiden, The Netherlands

10. Department of Neurology, Brain Center Rudolf Magnus, University Medical Center Utrecht, Utrecht, The Netherlands

11. Department of Epidemiology, ErasmusMC, Rotterdam, The Netherlands

12. Sequence Analysis Support Core, Leiden University Medical Center, Leiden, The Netherlands

13. SURFsara, Amsterdam, the Netherlands

14. Genomics Coordination Center, University Medical Center Groningen, University of Groningen, Groningen, the Netherlands

15. Medical Statistics Section, Department of Medical Statistics and Bioinformatics, Leiden University Medical Center, Leiden, The Netherlands
